# Evaluation of full-length *16S* rRNA amplicon sequencing using Oxford Nanopore Technologies for diversity surveys of understudied microbiomes

**DOI:** 10.1101/2025.09.20.676612

**Authors:** Vivian Yifan Li, Raúl González Pech, Diego Lera-Lozano, Verónica Román-Reyna, Mónica Medina

**Affiliations:** Department of Ecology and Evolutionary Biology, University of California Los Angeles, Los Angeles, CA, USA; Department of Biology, Texas State University, San Marcos, TX, USA; Department of Biology, The Pennsylvania State University, University Park, PA, USA; Department of Plant Pathology and Environmental Microbiology, The Pennsylvania State University, University Park, PA, USA

## Abstract

1. The use of long-read sequencing using portable Oxford Nanopore Technologies (ONT) is becoming increasingly popular in the study of host-associated microbiomes. However, its application has not yet been optimized for characterizing understudied microbiomes, such as those in marine environments.
2. We evaluated the accuracy and consistency of ONT sequencing of full-length *16S* rRNA genes for diversity surveys of symbiotic dinoflagellate (family Symbiodiniaceae) microbiomes.
3. When comparing amplicon sequencing of the full-length bacterial *16S* rRNA gene with only its V4 hypervariable gene region using a known microbial community, the former recapitulated the bacterial taxonomic composition more accurately.
4. ONT sequencing was also highly consistent between sequencing runs and flow cells.
5. Long-read sequencing technologies enable microbiome surveys using the full-length *16S* rRNA gene, achieving higher accuracy and resolution.
6. This work validates ONT long-read sequencing as a powerful tool for marine microbiome studies to catalyze advancements in the fields of ecology and evolution, resource management, and conservation.

## 1 INTRODUCTION

DNA sequencing has facilitated significant improvements in our understanding of complex marine holobionts through molecular characterization (González-Pech et al., 2024). For example, genetic markers have revealed greater divergence and genetic diversity than previously assumed within endosymbiotic dinoflagellates, leading to a revision of their systematics from the single genus *Symbiodinium* to the family Symbiodiniaceae, this latter comprising multiple genera (LaJeunesse et al., 2018). High-throughput sequencing technologies have further uncovered cnidarian microbiomes (McCauley et al., 2023; Planes et al., 2019; Pollock et al., 2018), prompting new discoveries about the role of bacteria in coral health and fitness amidst rapidly changing environments (Epstein et al., 2025; Roitman et al., 2020; Rosales et al., 2023).

The composition of marine microbiomes is commonly characterized by short-read shotgun sequencing of the bacterial small subunit ribosomal RNA (*16S* rRNA) gene. Though this gene is approximately 1,500 bp, commonly used short-read sequencing technologies (e.g., Illumina MiSeq) can only produce reads up to 300 bp (Stevens et al., 2023). It is thus a common practice to sequence one or two hypervariable regions of *16S* rather than the full gene. For instance, the Earth Microbiome Project (EMP) and the Global Coral Microbiome Project (GCMP) employ widely standardized protocols that target the V4 region (Epstein et al., 2019; Greg Caporaso et al., 2023; Pollock et al., 2018).

The hypervariable regions of *16S* rRNA vary in discriminatory power and detection biases, generating significant differences in the microbiome profiles determined by different regions (Hrovat et al., 2024; Lin & Ju, 2023). In contrast, full-length *16S* sequencing more accurately recapitulates known bacterial community profiles (Hrovat et al., 2024). Long-read sequencing, such as that offered by Oxford Nanopore Technologies (ONT), enables bacterial classification using the full *16S* rRNA gene, thus circumventing caveats associated with short reads. The ONT MinION platforms are also highly portable, which shortens the turnaround time between sample collection in the field and the implementation of actionable outcomes while simplifying the logistics of sample transportation and storage (Ames et al., 2021,Boykin et al., 2019, Carradec et al., 2020).

The application of ONT sequencing has not yet been optimized for understudied microbiomes, such as those in marine environments. Before ONT can be confidently used for future studies, an evaluation of its accuracy and consistency is required. A better understanding of sequencing yield and potential batch effects between flow cells and sequencing runs will also inform future experimental designs.

In this study, we assessed the reliability of ONT long-read sequencing to characterize microbiome composition by comparing the results of full-length bacterial *16S* rRNA sequencing (~1,500 bp) with sequencing of the V4 hypervariable gene region (~350 bp) using a known microbial community as reference. In addition, we used ONT to survey six Symbiodiniaceae cultures harboring distinct bacterial microbiomes to assess consistency between sequencing runs and to determine optimal read depth. We further tested this method in the field to assess the impact of inevitable environmental contamination when working in minimally equipped field stations. Our results demonstrate that long-read ONT sequencing of the full-length *16S* rRNA gene is highly accurate and consistent. It also achieves fine taxonomic resolution with sufficient read depth to fully capture species richness. This work validates and informs future use of ONT for understudied microbiomes.

## 2 MATERIALS AND METHODS

### 2.1 Study cultures

Symbiodiniaceae cultures were obtained from: the LaJeunesse Lab/Trench collection (The Pennsylvania State University, USA), the Medina Lab (The Pennsylvania State University, USA), and the Functional Phycology Lab (University of Aveiro, Portugal). Additional information about culture IDs, geographical origin, and host can be found in Supplementary Table 1. Culture maintenance, preparation and taxonomic verification methods are detailed in the Supporting Information. Experimental cultures (six strains, three replicates each) were incubated in ASP-8A medium for three weeks until late exponential phase of growth along with negative controls (three replicates) containing only ASP-8A medium without inoculation with Symbiodiniaceae.

### 2.2 Microbiome sampling

Microbiome samples from each Symbiodiniaceae culture, as well as negative controls, were collected at about the same time of day (11 am to 1 pm) to avoid differences attributed to diel microbiome cycling in these dinoflagellate cultures (Maire et al., 2021). Only bacteria that were physically attached to the Symbiodiniaceae cell surface were sampled following Maire et al. (2021) under sterile conditions as described in the Supporting Information.

### 2.3 DNA extraction

DNA extraction was performed using the DNeasy PowerSoil Pro kit (Qiagen) according to the manufacturer’s protocol. Two replicates of a mock community, the ZymoBIOMICSMicrobial Community Standard (Zymo Research), were included as positive controls to check the accuracy of library preparation, sequencing, and data processing. DNA extracts were stored at –20 °C until further use. DNA extracts were subsequently used for both Symbiodiniaceae genotyping with the eukaryotic large subunit ribosomal RNA gene (*28S* rRNA; see Supporting Information) and for PCR amplification of the full-length *16S* rRNA gene for microbiome characterization.

### 2.4 Short-read *16S* amplicon sequencing and processing

The full sequencing and processing procedure for short-read bacterial *16S* rRNA amplicons is described in the Supporting Information. Briefly, the V4 region of the *16S* rRNA gene was amplified in 20 µl PCR reactions following the Global Coral Microbiome Project (GCMP) amplicon sequencing protocol using the 515F/806R primer pair (Epstein et al., 2025) with Illumina IDT linkers attached to the 5’ end (Supplementary Table 2). A full DNA extraction and amplification procedure is described in the Supporting Information. A sample of autoclaved nuclease-free water was included as a negative PCR control. PCR products were submitted to the Genomics and Microbiome Core Facility at Rush University Medical Center for library preparation, paired-end sequencing on an Illumina MiSeq i100+, and sequence processing.

Sequences received from the sequencing facility were processed and analyzed in R v4.4.1. Taxonomy was assigned using DADA2 v1.35.0 (Callahan et al., 2016) and the SILVA *SSU* database version 138. Assignments with bootstrap confidence values <80% were classified as “NA”. Sequences that mapped to mitochondrial or chloroplast *16S* rRNA genes were removed using phyloseq v1.48.0 (McMurdie & Holmes, 2013).

### 2.5 Full-length *16S* amplicon sequencing and processing

The complete sequencing and processing procedures for full-length bacterial *16S* amplicons are described in the Supporting Information. Briefly, the *16S* rRNA gene was amplified in 25 µl PCR reactions using the degenerate *16S* rDNA primer set as described in Waechter et al. (2023): S-D-Bact-0008-c-S-20/S-D-Bact-1492-a-A-22 with anchor sequences attached to the 3’ end (Supplementary Table 2, Klindworth et al., 2013; Matsuo et al., 2021). A sample of autoclaved nuclease-free water was included as a PCR negative control. Due to low yield of bacterial DNA extracts and thus PCR products, each PCR reaction was replicated six times and then pooled between all six replicates, then purified using the DNA Clean & Concentrator-5 PCR purification kit (Zymo Research) and AMPure XP beads (Agencourt AMPURE XP, Beckman Coulter).

A multiplexed library (20 barcodes) was prepared using the Native Barcoding Kit 24 V14 (SQK-NBD114.24) and loaded onto separate R10.4.1 flow cells (FLO-MIN114). The same library was sequenced three times on different flow cells of varying health (Table 1) that had been previously used and washed following the manufacturer’s instructions. This was done to determine the appropriate sequencing depth and the variability of sequencing output produced by separate sequencing runs. Sequencing was performed on a MinION Mk1C controlled by MinKNOW v 23.07.12 on an iMac (High Sierra Version 10.13.6, Intel Core i7, 32GB, 4 cores) connected by USB, with real-time basecalling.

**Table 1.** Duration of each sequencing run and flow cell health (i.e., number of active pores) before and after sequencing.

| Run ID | Run duration | Flow cell health before sequencing | Flow cell health after sequencing |
| --- | --- | --- | --- |
| Run 1 | 72 h | 1,029 | ~1 |
| Run 2 | 60 h | 764 | ~2 |
| Run 3 | 40 h | 237 | 0 |

Long-read processing was conducted on the Roar Collab High Performance Computing Cluster at The Pennsylvania State University (code available at: https://github.com/VivianYifanLi/Li_et_al_2025_ONTSequencing). Method details are available in the Supporting Information. Additional basecalling, demultiplexing, and adaptor and barcode trimming were done using Dorado v0.5.3 (ONT). Dorado-basecalled sequences were used for all downstream analyses. The ONT EPI2ME platform (*wf-16S*) was used for alignment and taxonomic assignment to reference sequences from the NCBI *16S* + *18S* rRNA and SILVA v138 SSU (small subunit) databases. Alignment was done using minimap2 v2.29 (Li, 2021) followed by taxonomic assignment based on a percentage identity of ≥99% for NCBI (species level) and ≥95% for SILVA sequences (genus level), respectively. A lower percentage identity cut-off was used for SILVA as the database does not curate species-level taxonomy. Amplicons with lower percentage identities were classified as “Unknown” and their proportions determined through manual inspection in R v 4.4.1 using phyloseq v1.48.0 (McMurdie & Holmes, 2013).

### 2.6 Microbiome analysis, consistency between sequencing runs, and optimal sequencing depth

DADA2 (short reads) generated an Amplicon Sequences Variant (ASV) table while EPI2ME (long reads) generated a taxonomic abundance table down to genus and species respectively from SILVA and NCBI. Both tables were imported into R for downstream analyses using phyloseq v1.48.0 (McMurdie & Holmes, 2013). Taxa with <1% abundance and <10 reads (the number of reads in the negative controls) per sample were filtered from all samples. Decontam v1.24.0 (Davis et al., 2018) was used to identify and remove contaminants in the data using the combined approach (Supplementary Figure 1).

Relative abundances were center log-ratio transformed using the R package compositions v2.0-8 (van den Boogaart et al., 2025). A pseudocount of 1 was added to all OTU counts to enable log-ratio transformation of zero values. Principle coordinate analysis (PCoA) and non-metric dimensional scaling (NMDS) plots were generated using ggplot2 v 3.5.2 (Wickham, 2016) and used to visualize community dissimilarities based on Aitchison distances using phyloseq v1.48.0. Analysis of similarities (ANOSIM) and Permutational Multivariate Analysis of Variance (PERMANOVA) were performed to test for significant differences between sequencing runs using the R packages vegan v2.7.1 (Oksanen et al., 2025), veganEx v0.1.0 (*GitHub - ZhonghuiGai/VeganEx*, 2021) and RVAideMemoire v0.9-83-7 (HERVE, 2025), with 999 permutations per test. Venn diagrams were plotted using ggVennDiagram v1.5.2 (Gao et al., 2024).

To estimate the optimal sequencing depth for our samples, rarefaction curves with extrapolation were plotted using the R packages iNEXT v3.0.1 (Hsieh et al., 2025) and grid v4.4.1 from the sequencing run with the most data. To determine whether the microbial composition determined by our sequencing approaches acurately recapitulated the actual microbial composition within each sample, the Good’s coverage index was calculated using the equation 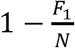 where *F*_1_ is the number of singletons and *N* is the total number of individuals in each sample. A higher Good’s coverage index indicates a higher probability that all taxa present in the sample have been detected.

### 2.13 Field testing

All procedures described next were performed in the Key Largo Marine Research Laboratory (KLMRL; a non-sterile field station) in Key Largo, Florida. DNA was extracted, amplified and purified following the procedure described above and in the Supporting Information from three replicates of the ZymoBIOMICS Microbial Community Standard (75 µl each) along with a negative control of nuclease-free water (1 ml). DNA amplification was performed only once with 30 amplification cycles instead of six repetitions with 25 amplification cycles. Concentrations of PCR products were quantified using a Quantus fluorometer (Promega) with ONE dsDNA system. To expedite the procedure in the field, library preparation and metabarcoding was performed using the Rapid Barcoding Kit 24 V14 (SQK-RBK114.24) following the ONT protocol “Rapid sequencing V14 – Amplicon sequencing”.

The multiplexed library was sequenced on an R10.4.1 flow cell (FLO-MIN114) on the MinION Mk1B monitored by MinKNOW v24.11.10 on a MacBook Pro (Sonoma v14.6.1, Apple M1, 16GB, 8 cores), without real-time basecalling. Dorado basecalling, sequence processing and bioinformatics analyses were performed following procedures described above and in the Supporting Information.

## 3 RESULTS

### 3.1 Short-read sequencing generated more reads, but long-read sequencing achieved finer taxonomic resolution

Nanopore MinION (ONT) sequencing generated 2,485 to 31,514 reads per sample across three sequencing runs (18 samples each) after singleton removal and decontamination (Figure 1; Supplementary Table 3). A total of 75 unique genera as determined by alignment to the SILVA v138 *SSU* database were detected in every sequencing run. Illumina MiSeq sequencing generated 3,259 to 55,136 reads per sample after singleton removal and decontamination. A total of 20 unique genera were detected.

**Figure 1.**
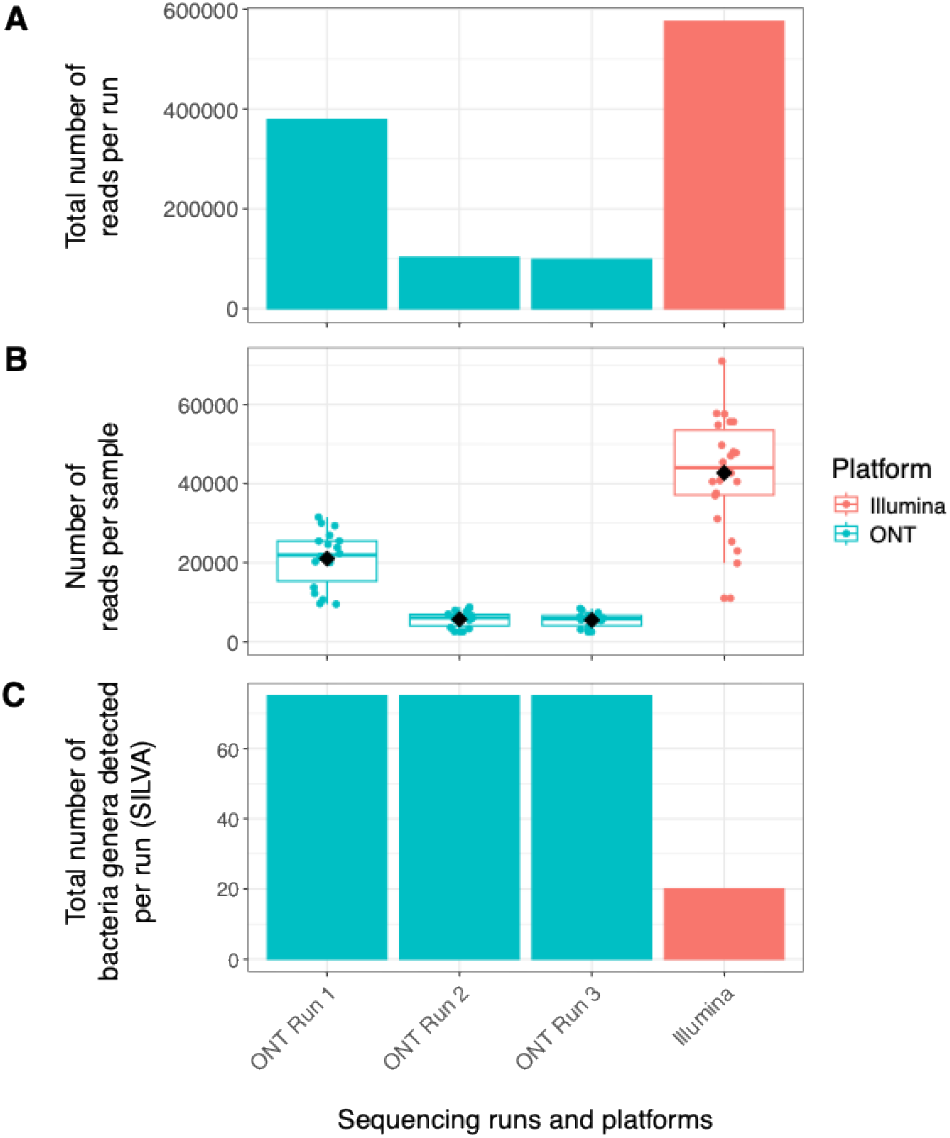
Summary of (A) total sequencing yield, (B) yield per sample, and (C) number of bacterial genera detected across all ONT (n=3) and Illumina (n=1) sequencing runs. Black diamonds in (B) represent the mean number of reads across samples per sequencing run.

Four genera (*Granulicatella, Filomicrobium, Thalassobaculum* and an uncultured Flavobacteriaceae) were found by decontam v1.24.0 to be results of putative contaminant ONT sequences and removed before further analyses. In contrast, 41 amplicon sequence variants (ASVs) corresponding to two genera (*Pyruvatibacter* and an unclassified Saprospiraceae) were detected as putative contaminants in the Illumina dataset and removed.

### 3.2 Species-level taxonomic profiling of a mock bacterial community

Species-level taxonomic profiling was only done with the ONT data. Full-length *16S* amplicon sequencing correctly identified the presence of five out of the eight species in the mock bacterial community when using the NCBI *16S*/*18S* database as reference, with the exceptions of *Bacillus subtilis, Escherichia coli* and *Listeria monocytogenes* (Supplemental Figure 2). Additionally, 12 species were misclassified at the species level. Sequences classified as “Unknown” made up roughly 28-32% of all sequences across the three sequencing runs.

### 3.3 Genus-level taxonomic profiling of a mock bacterial community

Taxonomic classification of full-length *16S* rRNA amplicons at the genus level was more accurate than at the species level. The SILVA 138 SSU and NCBI *16S*/*18S* databases performed similarly (Supplemental Figure 3). At the genus level, long-read sequencing correctly identified the presence and relative abundances of all eight genera in the mock community when using the SILVA database (Figure 2A, B). Short-read sequencing targeting the V4 hypervariable region failed to identify the presence of two genera: *Pseudomonas* and *Salmonella* (Figure 2A, D). The proportion of unclassified or unknown taxa was also higher compared to full-length sequencing.

**Figure 2.**
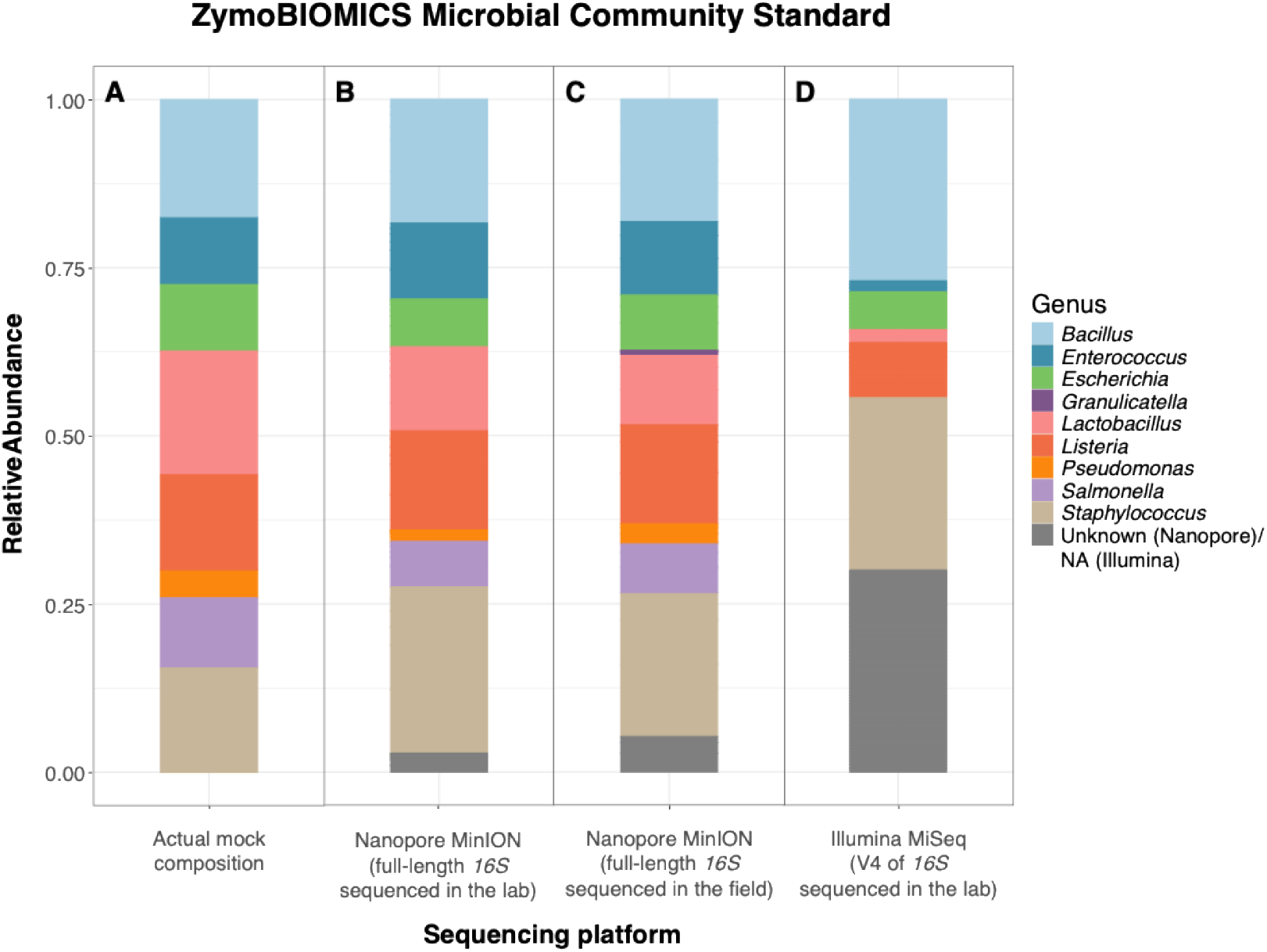
Comparison of the actual mock composition (A) and observed relative abundances of bacterial genera found within the ZymoBIOMICS Microbial Community Standard, based on full-length 16S sequencing in the lab (B) and the in the field (C), and short-read sequencing of the 16S V4 region in the lab (D). Expected composition of the mock community was provided by the manufacturer (Zymo Research, 2023). Composition of the Zymo Microbial Community Standard was consistent between sequencing runs and replicates, thus all replicates were pooled in these relative abundance plots.

### 3.4 Field testing

ONT library preparation and sequencing performed in the field yielded similar results to those performed in the lab (Figure 2). However, in the field, *Granulicatella* was not identified and removed as a contaminant by the decontamination step when the same classification thresholds were used (Supplementary Figure 1).

Overall, long-read sequencing generated data that was more reliable than that of short-reads in both lab and field settings. Our results will focus on the full-length amplicon sequencing approach with genus-level taxonomy referencing the SILVA database.

### 3.5 No batch effects detected across full-length *16S* sequencing runs

Results were highly consistent between three separate full-length *16S* sequencing runs even though flow cells of different conditions (Table 1) were used for each run (Figure 3). There were no significant differences between group centroids (PERMANOVA p=0.85, 999 permutations) and within-group similarities were greater than between group similarities (ANOSIM p=0.347, 999 permutations). All 75 detected genera were shared between all three sequencing runs (Figure 3C), i.e., no genus was unique to any run.

**Figure 3.**
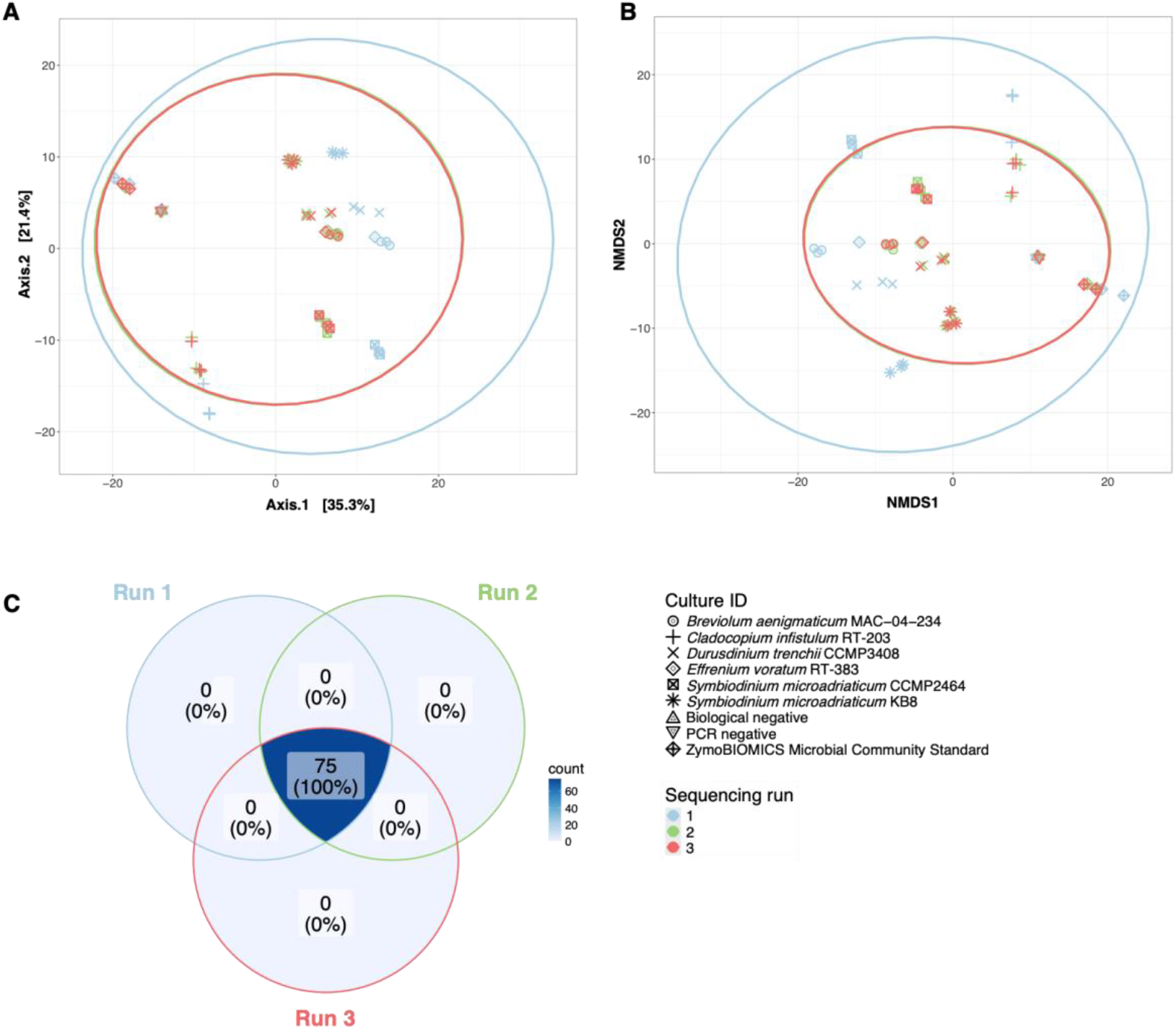
Consistency of the observed microbiome composition of six Symbiodiniaceae cultures as determined by three full-length 16S sequencing runs on a Mk1C sequencing device using different flow cells. (A) PCoA and (B) NMDS plots (stress=0.1132943 show the Aitchinson distances between results from three separate sequencing runs. (A-B) Ellipses represent 95% confidence intervals. (C) All 75 detected genera were shared between all three sequencing runs.

### 3.6 Species richness in Symbiodiniaceae culture microbiomes is sufficiently captured in 9,525 to 31,554 reads

Data from Run 1 of full-length *16S* rRNA sequencing was used to estimate optimal sequencing depths. Each sample contained 9,525 to 31,554 reads. The Good’s coverage indices suggested that >99% of taxa (Figure 4A) were captured in all samples. Extrapolated rarefaction curves suggested that 20,000 reads sufficiently capture the species richness in all samples (Figure 4B).

**Figure 4.**
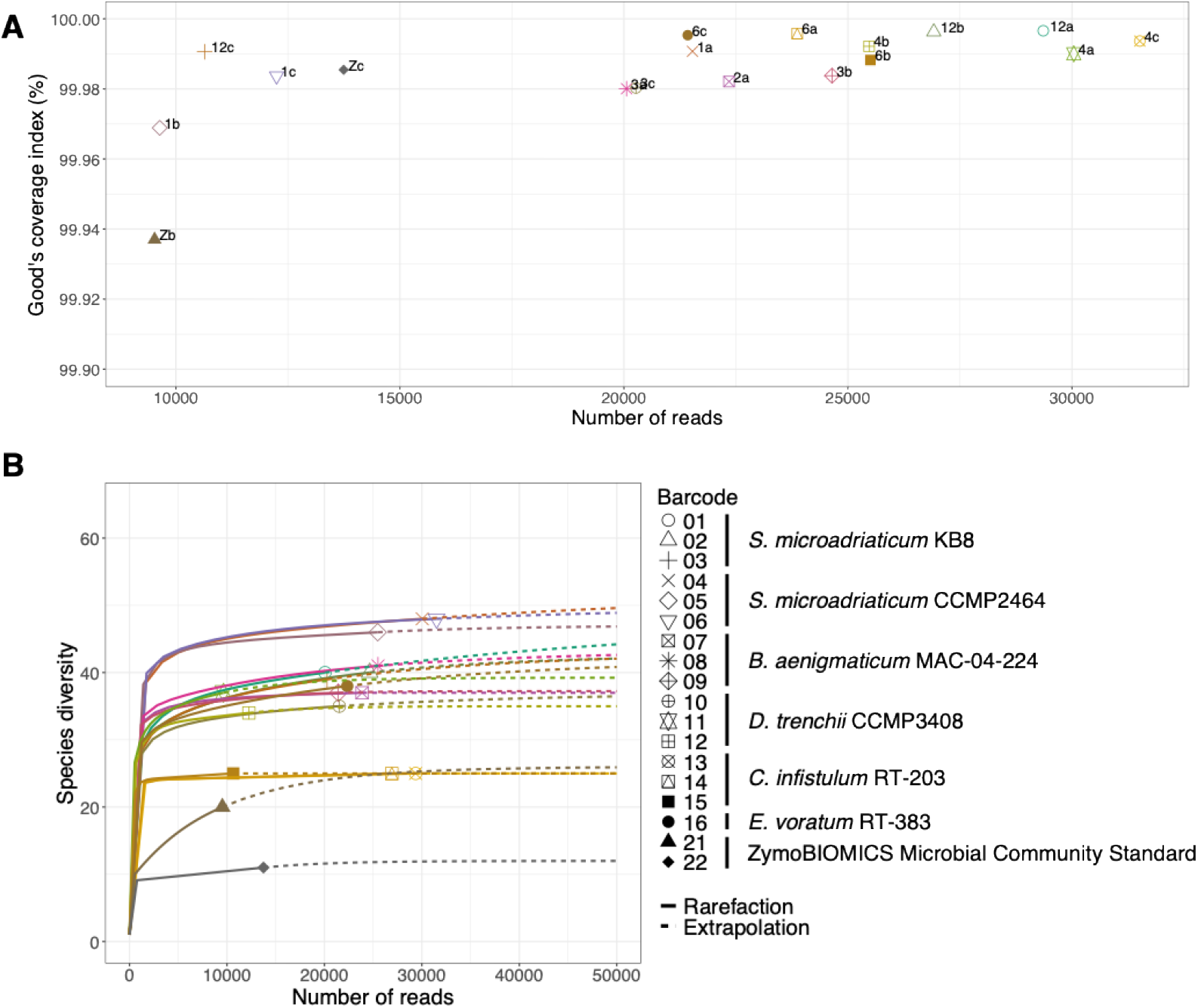
Optimal read depth determination using data from full-length 16S sequencing Run 1. (A) Good’s coverage index per sample plotted against numbers of reads per sample. Each color and symbol represent a barcode included in the multiplexed library and each barcode is labelled with its corresponding sample ID. (B) Rarefaction curves where each color and symbol represent a barcode included in the multiplexed library. Solid lines represent observed data while dashed lines represent data extrapolated to 50,000 reads.

### 3.7 Symbiodiniaceae microbiomes exhibit both inter- and intraspecific variation

Symbiodiniaceae cultures harbor distinct microbiomes (ANOSIM p=0.001; PERMANOVA p=0.001), with differences between species but also between strains within a same species (*e*.*g*.. *Symbiodinium microadriaticum* KB8 and CCMP2464; ANOSIM p_adj_=0.015; PERMANOVA p_adj_=0.0015; Figure 5). There were 11 bacterial taxa shared between all cultures, including *Labrenzia* and *Marinobacter*.

**Figure 5.**
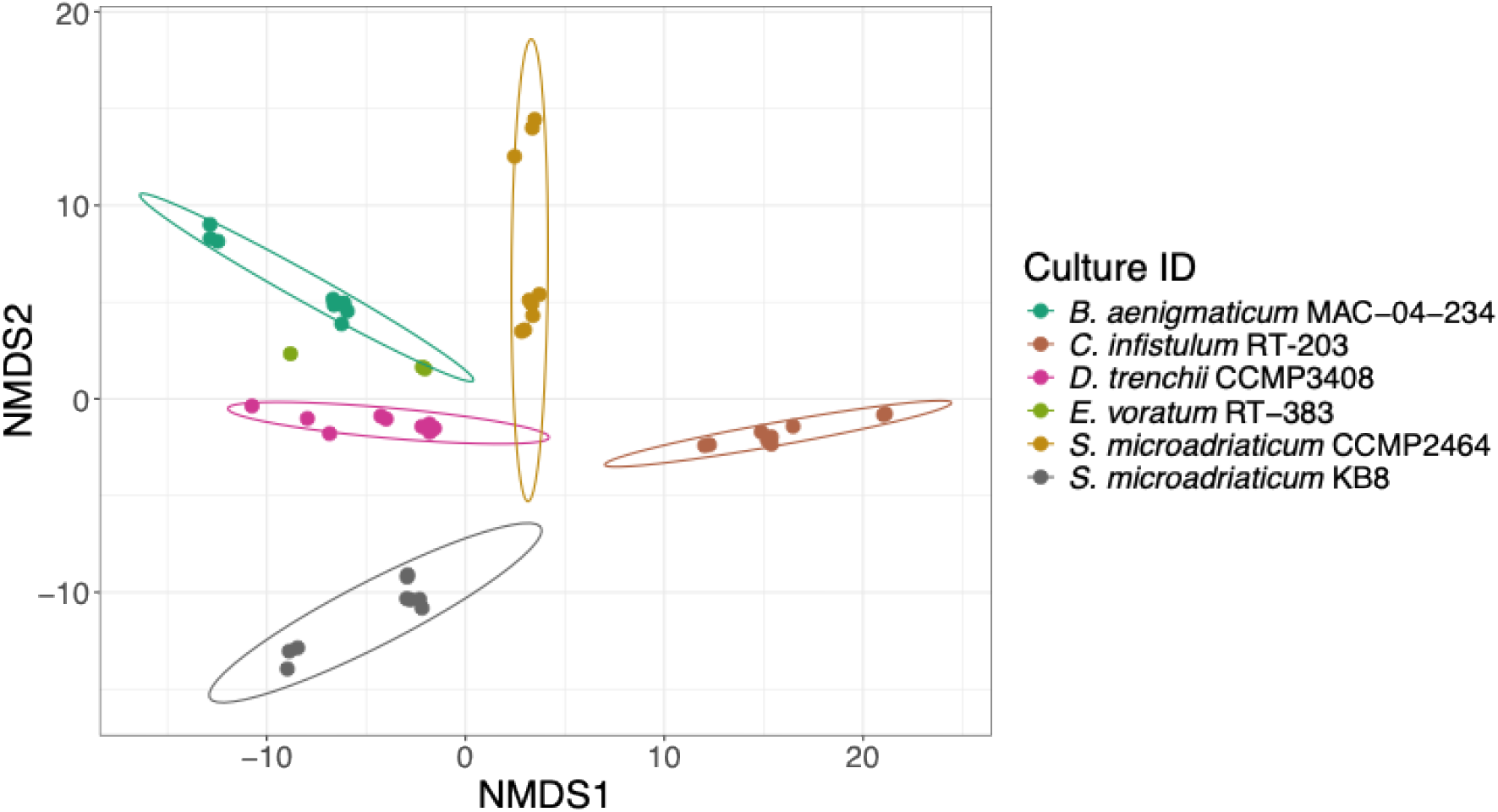
Bacterial microbiomes of six Symbiodiniaceae cultures form distinct clusters on an NMDS plot (stress=0.1148755) based on Aitchison distances between the six cultures. Ellipses represent 95% confidence intervals for cultures with at least n = 5 after all sequencing runs were pooled.

## 4 DISCUSSION

Long-read sequencing platforms such as Oxford Nanopore Technologies are becoming more accessible, enabling microbiome researchers to overcome the traditional constraints of short-read *16S* rRNA gene sequencing. However, there has been a lack of standardization and validation of such methods for exploring understudied environments such as marine holobionts. In this study, we benchmarked the accuracy and consistency of full-length bacterial *16S* rRNA gene sequencing using ONT against a short-read sequencing pipeline used for the Global Coral Microbiome Project (Pollock et al., 2018). Full-length *16S* rRNA sequencing was also used to describe inter- and intra-specific specific variation in Symbiodiniaceae microbiomes as a case study. Long-read sequencing generated results that were more accurate down to the genus level than short-read sequencing in its survey of bacterial microbiomes.

### 4.1 Long-read sequencing is highly accurate and consistent

Historically, ONT has been associated with high error rates and low sequencing depth (Carradec et al., 2020). In this study, we achieved high accuracy in our bacterial composition surveys despite these error rates. Long-read sequencing of the full length *16S* rRNA gene was more accurate than short-read sequencing in identifying the presence/absence and relative abundances of taxa within a Microbial Community Standard (Figure 2).

The MinION generated highly consistent results between sequencing runs regardless of flow cell health. While sequencing yield was low compared to that of the Illumina MiSeq, negligible batch effects in ONT sequencing enables samples to be sequenced across multiple runs and then analyzed together. Thus, fewer samples could be run at a time to increase sequencing yield.

Sequencing conducted in the field produced results comparable to those obtained in the laboratory. Nonetheless, the persistence of potential contaminant taxa even after decontamination indicates that decontam settings may need to be adjusted based on the specific datasets being analyzed.

### 4.2 Long-read sequencing identified core members of Symbiodiniaceae microbiomes consistent with previous studies

We used full-length *16S* amplicon sequencing to survey the microbiomes of six lab cultures of Symbiodiniaceae across different Symbiodiniaceae strains, species and genera. We extracted bacteria physically attached to the cell surface of Symbiodiniaceae based on the assumption that cells located in closer proximity to each other perform significant levels of nutrient exchange (Stevens et al., 2023). Across all three sequencing runs, we consistently detected a total of 75 bacterial genera across six Symbiodiniaceae cultures, with an average of 45 genera per culture.

Symbiodiniaceae cultures assessed in this study harbor distinct microbiomes but also host 11 bacterial genera that were shared between all Symbiodiniceae genera. Out of these 11 bacterial genera, *Labrenzia* and *Marinobacter* were also found to be core members of the Symbiodiniaceae microbiome in a previous study (Lawson et al., 2018) despite differences in isolation method, culture maintenance practices, growth media, and growth conditions.

### 4.3 Limitations of ONT long-read sequencing

The ONT pipeline described in this study depends on stringent percentage identity cut-offs and filtering of any taxa that cannot be assigned a genus, which may discard sequences that do not have good representative references in public databases. Such sequences are common in understudied environments such as marine invertebrate holobionts (González-Pech et al., 2024; Stévenne et al., 2021), which may lead to an underrepresentation of uncharacterized bacterial taxa. Nonetheless, this pipeline provides high levels of accuracy in identifying the taxa that can be identified. Percentage identity cut-offs can also be modified for studies with different requirements.

Low sequencing yield in this study is most likely the result of using old and/or used flow cells. In a different study exploring agricultural soil microbiomes, the MinION generated 1,695,436 total sequence reads from a single run, averaging 66,843 reads per sample (n=25, Stevens et al., 2023). This is comparable to the number of reads we generated from short-read sequencing on the Illumina MiSeq. Since this study examined only bacteria that were physically attached to the surface of a small number of Symbiodiniaceae cells, the lower read depths may be better suited to our samples because of the low microbial biomass and lower community diversity compared to more complex microbiomes. Optimal sequencing depths for other sample types and other microbiomes may thus vary.

### 4.4 Differences in taxonomic assignment algorithms between long and short reads may introduce biases

The DADA2 approach to sequence processing, denoising and taxonomic classification via amplicon sequence variants (ASVs) depends on the presence of a substantial proportion of reads (>10% of reads) in each sample that contain zero errors (Callahan, 2022). Long reads of the full length *16S* rRNA gene may not consistently meet this assumption (Callahan, 2022). The core denoising algorithm in DADA2 was built based on error rates in Illumina-sequenced reads (Callahan et al., 2016) Therefore, DADA2 was only used for processing short reads while EPI2ME/*wf-16s* was used for long reads in this study.

DADA2 and EPI2ME/*wf-16s* utilize different algorithms that could introduce biases that are important to consider for future applications. DADA2 bins highly similar reads into partitions, each of which represents an amplicon sequence variant (ASV) (Callahan et al., 2016). Taxonomies are then assigned to each ASV, rather than to individual reads, using the RDP Naive Bayesian Classifier algorithm that is based on *k*-mers. In contrast, EPI2ME/*wf-16s* assigns a taxonomy to every sequenced read by aligning every whole read to a reference database and calculating the percentage identity of its highest scoring alignment using minimap2 (Oxford Nanopore Technologies, n.d.). Thus, DADA2 may be more effective at correcting for sequencing errors in individual reads. However, true sequence variants could sometimes be misconstrued as erroneous and thus assigned to an incorrect partition (Callahan et al., 2016). By comparison, EPI2ME/*wf-16s* classifies all reads equally, which better detects individual sequence variants but does not differentiate erroneous reads from true sequence variants without the use of additional tools.

## 5 CONCLUSION

Full-length *16S* rRNA gene sequencing enables characterization of microbiome members at high accuracy, consistency, and fine taxonomic resolution. We recommend this method for sample sizes of up to 24, which can be multiplexed into a single library, and for field-based studies that depend on quick turnaround times. The ONT MinION platforms are relatively low-throughput and thus inefficient for large sample sizes containing highly diverse microbiomes such as those in the GCMP. For numerous, highly diverse samples, one could consider the ONT GridION or PromethION to retain the benefits of long-read sequencing while increasing sequencing yield.

Long-read sequencing technologies are becoming increasingly competitive with short-read sequencing technologies in terms of accuracy and yield while providing the added capability of sequencing larger gene fragments, thus improving the taxonomic reliability of *16S* surveys. This holds exciting promise for future studies of understudied microbiomes that could catalyze advancements in the fields of ecology and evolution, resource management, and conservation.

## ACKNOWLEDGEMENTS

This material is based upon work supported by the National Science Foundation under Award No. 2227070. VYL was partially supported by the Verne M. Willaman Distinguished Graduate Fellowship in Science from the J. Jeffrey and Ann Marie Fox Graduate School at The Pennsylvania State University. We would like to thank Todd LaJeunesse, Jörg Frommlet, and Anthony Bellantuono for providing Symbiodiniaceae cultures that allowed us to conduct this research. We are grateful to Dr. William Fitt and the University of Georgia for hosting us at the Key Largo Marine Research Station (KLMRL) during the field portion of this study, as well as the students in Penn State’s BIOL 483: Coastal Biology Field Experience course for their help with sample collection at KLMRL. Additionally, we would like to thank Elena Bollati and Alessandro Garritano for bioinformatics assistance. Finally, we are grateful to Todd LaJeuneese and Gui Becker for their constructive comments.

## DATA AVAILABILITY

Raw sequencing data are available under the BioProject accession number xxxxx.

## AUTHOR CONTRIBUTION

VYL, RGP and MM conceptualized the project and designed the experiments. VYL performed the experiments and collected the data. VYL and DLL processed the raw sequencing data. VYL, RGP and VRR participated in designing and running data analysis. VYL wrote the manuscript and all authors edited, reviewed and approved the manuscript.

