## Supporting Information for "Evaluation of full-length *16S* rRNA amplicon sequencing using Oxford Nanopore Technologies for diversity surveys of understudied microbiomes"

#### Symbiodiniaceae culture maintenance and preparation of experimental cultures

Cultures had been maintained in either F/2 (LaJeunesse Lab) or ASP-8A media (Medina and Functional Phycology Labs), with transfers into fresh medium every two to four months. At the onset of this experiment, 0.5 ml of each experimental culture (six strains, three replicates each) were transferred into 4.5 ml of fresh, autoclaved ASP-8A medium using sterile technique. Cultures were incubated for three weeks until late exponential phase of growth under the following conditions: temperature of 27 °C, light intensity of 40-50 μmol quanta m-2 s-1, light regime of 12 h light and 12 h dark. Negative controls (three replicates) were prepared in the same manner using only ASP-8A medium without inoculation with Symbiodiniaceae.

The taxonomic identity of each Symbiodiniaceae culture was verified via Sanger sequencing of the eukaryotic *LSU* marker gene following LaJeunesse et al. (2018). Amplicons were directly sequenced on an Applied Biosciences sequencer (Applied Biosciences, Foster City, CA, USA) at The Pennsylvania State University Genomics Core Facility. Chromatograms were checked, and sequences aligned, using Geneious v9.1.6.

#### Microbiome sampling from Symbiodiniaceae cultures

Symbiodiniaceae cells in each culture were resuspended in the growth medium, 1 ml of resuspended cells were collected and filtered through individual 5 μm-mesh strainers (pluriSelect, USA). Strainers were sealed with parafilm and centrifuged for 5 min at 12,000 *g* to separate suspended bacteria (<5 μm) from Symbiodiniaceae cells (>5 μm). The same strainers were then rinsed once with 500 μl autoclaved ASP-8A media and centrifuged for another 5 min at 12,000 *g*. The filters, having retained Symbiodiniaceae cells and bacteria cells tightly attached to the dinoflagellate cell surface, were retrieved from the strainer using sterile forceps, placed into sterile 1.5 ml microcentrifuge tubes respectively, and stored at –80 °C until DNA extraction. Three blank strainers were rinsed once with 1 ml and a second time with 500 μl of sterile ASP-8A media, respectively, to serve as negative controls.

#### Short-read *16S* amplicon sequencing and processing

Following the Global Coral Microbiome Project (GCMP) amplicon sequencing protocol (Epstein et al., 2025; Pollock et al., 2018), the *16S* V4 region was amplified using the 515F/806R primer pair with Illumina IDT linkers (underlined) attached to the 5' end: 515F 5'-CTACACGACGCTCTTCCGATCTGTGYCAGCMGCCGCGGTAA-3' and 806R 5'-CAGACGTGTGCTCTTCCGATCTGGACTACNVGGGTWTCTAAT-3'. Each 20 μl PCR reaction contained 0.4 μl dNTPs (10 mM ), 0.6 μl $MgCl_2$ (50 mM), 2 μl buffer (10x concentrated), 0.1 μl Taq DNA polymerase (5 U $\mu l^{-1}$), 1 μl of forward primer (10μM), 1 μl of reverse primer (10 μM), and 13.9 μl of autoclaved nuclease-free water. A sample of autoclaved nuclease-free water was included as a negative control to check for any contamination in the PCR reagents. Reactions were held at 94 °C for 2 min for initial denaturation, amplified in 35 cycles at 94 °C for 45 s, 55 °C for 1 min and 68 °C for 90 s, then underwent a final extension at 68 °C for 7 min.

PCR products were submitted to the Genomics and Microbiome Core Facility at Rush University Medical Center for library preparation and paired-end sequencing on an Illumina MiSeq i100+ platform (maximum read length: 2 x 300 bp). Data preprocessing including primer and adaptor removal, trimming and filtering by sequence length, and filtering by sequence quality were performed by the facility.

Sequences received from the sequencing facility were processed and analyzed in R v4.4.1. DADA2 v1.35.0 (Callahan et al., 2016) was used to correct sequence errors, dereplicate sequences, merge forward and reverse overlapping reads, remove chimeras, infer amplicon sequence variants (ASVs), and assign taxonomy using the RDP Naive Bayesian Classifier algorithm (*k*-mer size 8, 100 bootstrap replicates) (Wang et al., 2007) and SILVA *SSU* database version 138. Assignments with bootstrap confidence values <80% were classified as "NA". Sequences that mapped to mitochondrial or chloroplast *16S* rRNA genes were removed using phyloseq v1.48.0 (McMurdie & Holmes, 2013).

**Full-length *16S* amplicon sequencing and processing**

The full-length bacterial *16S* gene was amplified using the degenerate *16S* rDNA primer set as described in Waechter et al. (2023): S-D-Bact-0008-c-S-20 with an anchor sequence (underlined) 5′-TTTCTGTTGGTGCTGATATTGCAGRGTTYGATYMTGGCTCAG-3′ and S-D-Bact-1492-a-A-22 with an anchor sequence (underlined) 5′-ACTTGCCTGTCGCTCTATCTTCCGGYTACCTTGTTACGACTT-3′ (Klindworth et al., 2013; Matsuo et al., 2021). Each 25 μl PCR reaction contained 2 μl of DNA template, 0.5 μl of forward primer (10 μM), 0.5 μl of reverse primer (10 μM), 12.5 μl of LongAMP Taq 2x Master Mix (New England Biolabs, USA), and 9.5 μl of autoclaved nuclease-free water. A sample of autoclaved nuclease-free water was included as a negative control to check for any contamination in the PCR reagents. Reactions were held at 95 °C for 1 min for initial denaturation, amplified in 25 cycles at 95 °C for 20 s, 51 °C for 30 s and 65 °C for 2 min, then underwent a final extension at 65 °C for 5 min. Due to low yield of bacterial DNA extracts and thus PCR products, each PCR reaction was replicated six times and then pooled between all six replicates.

Purification and concentration of the PCR products were performed using the DNA Clean & Concentrator-5 PCR purification kit (Zymo Research) followed by a second purification step with AMPure XP beads (Agencourt AMPURE XP, Beckman Coulter) on a magnetic bead separation rack (Bel-ArtTM SP Scienceware), at ambient temperature using a 1–1.6x ratio (to remove fragments <100 bp). Concentrations of PCR products were quantified using a Quantus fluorometer (Promega) with ONE dsDNA system. DNA purity based on A260/280 and A260/230 ratios were measured using a NanoDrop 2000c Spectrophotometer (Thermo Fisher Scientific). Library preparation and metabarcoding was done using the Native Barcoding Kit 24 V14 (SQK-NBD114.24) according to the ONT Early Access protocol "Ligation sequencing amplicons - Native Barcoding Kit 24 V14 (SQK-NBD114.24)", which enables multiplexing of up to 24 samples with unique barcodes.

A 15 fmol aliquot of the multiplexed library was loaded onto separate R10.4.1 flow cells (FLO-MIN114) and sequenced using the MinION Mk1C controlled by MinKNOW v23.07.12 on an iMac (High Sierra Version 10.13.6, Intel Core i7, 32GB, 4 cores) connected by USB. Live basecalling of raw reads was done using Guppy v7.1.4 (ONT) with a quality score cutoff of 8.

Long-read sequence processing was conducted on the Roar Collab High Performance Computing Cluster at The Pennsylvania State University (code available at: https://github.com/VivianYifanLi/Li_et_al_2025_ONTSequencing). Additional basecalling (implementing the simplex basecalling model dna_r10.4.1_e8.2_400bps_sup@v4.3.0), demultiplexing, and adaptor and barcode trimming for reads with quality scores above 8 as determined by real-time basecalling were done using Dorado v0.5.3 (ONT). Dorado-basecalled sequences were used for all downstream analyses. Cutadapt v4.0 was used to remove PCR primers. Prowler (Lee et al., 2021) was used to trim low-quality sequence windows (size=60 bp, static algorithm) with average per-base quality scores that fall below 20. Reads <200 bp were discarded.

The ONT EPI2ME platform was used for downstream sequence analysis: *wf-alignment* was used to check for errors in library preparation; *wf-16S* to remove reads <800 bp or >2000 bp and with average read quality <20. Subsequently, *wf-16S* was used for alignment and taxonomic assignment of full-length *16S* sequences to reference sequences from the NCBI *16S* + *18S* rRNA and SILVA v138 *SSU* (small subunit) databases. Alignment was done using minimap2 v2.29 (Li, 2021) followed by taxonomic assignment based on a percentage identity ≥99% for NCBI (species level) and ≥95% for SILVA sequences (genus level), respectively. A lower percentage identity cut-off was used for SILVA because the database does not curate species-level taxonomy. Amplicons with lower percentage identities were classified as "Unknown" and their proportions determined through manual inspection in R v4.4.1 using phyloseq v1.48.0 (McMurdie & Holmes, 2013).

### SUPPLEMENTARY MATERIALS

Supplementary Table 1. Species names and strain IDs of Symbiodiniaceae cultures used for microbiome characterization along with the culture collection they were obtained from, geographical origin, and host from which they were originally isolated.

| **Sample ID** | **Genus** | **Species** | **Strain ID** | **Available in collection** | **Original collection** | **Host species** | **Geographical origin** | **ITS Subtype** |
|---|---|---|---|---|---|---|---|---|
| 6 | *Breviolum* | *B. aenigmaticum* | MAC-04-225 | LaJeunesse | Coffroth | *Porites asteroides* | Caribbean | B1 |
| 3 | *Symbiodinium* | *S. microadriaticum* | KB8 | Medina | Coffroth | *Cassiopea xamachana* | North Pacific | A1 |
| 4 | *Symbiodinium* | *S. microadriaticum* | CCMP2464 | Medina | Trench | *Cassiopea xamachana* | North Atlantic | A1 |
| 2 | *Effrenium* | *E. voratum* | RT-383 | Medina | LaJeunesse | *Anthopleura elegantissima* | West Pacific | E1 |
| 1 | *Durusdinium* | *D. trenchii* | CCMP3408 | Medina | Coffroth | *Montastraea faveolata* | Caribbean | D1a |
| 12 | *Cladocopium* | *C. infistulum* | RT-203 | Functional Phycology Lab | Trench | *Hippopus sp.* | West Pacific | C2 |

Supplementary Table 2. Sequences of primer pairs used in PCR for short-read and long-read sequencing, respectively. Illumina IDT linkers and ONT anchor sequences are underlined.

| Platform | Read length | Primer pair | Primer sequence |
|---|---|---|---|
| Illumina | Short | 515F<br>806R | 5’-CTACACGACGCTCTTCCGATCTGTGYCAGCMGCCGCGGTAA-3'<br>5’-CAGACGTGTGCTCTTCCGATCTGGACTACNVGGGTWTCTAAT-3' |
| ONT | Long | S-D-Bact-0008-c-S-20<br>S-D-Bact-1492-a-A-22 | 5′-TTTCTGTTGGTGCTGATATTGCAGRGTTYGATYMTGGCTCAG-3′<br>5′-ACTTGCCTGTCGCTCTATCTTCCGGYTACCTTGTTACGACTT-3′ |

Supplementary Figure 1. Distribution of decontam score statistics P generated from NCBI species assignments (A) and SILVA genus assignments (B) from full length *16S* sequences, and SILVA genus assignments from V4 region sequences (C). The combined approach of decontam was used, which evaluates contaminants based on whether the sequence frequency varies inversely with sample DNA concentration and whether they have increased prevalence in negative controls (Davis et al., 2018). Low P scores indicate the taxa better fits the model that identifies it as a contaminant while high scores indicate the taxa better fits the model that identifies it as a non-contaminant. Bimodality was observed between very high and very low scores. A classification threshold $P^*$=0.5 was determined by identifying the decontam score cut-off that best identifies the contaminants that make up the low-score mode.

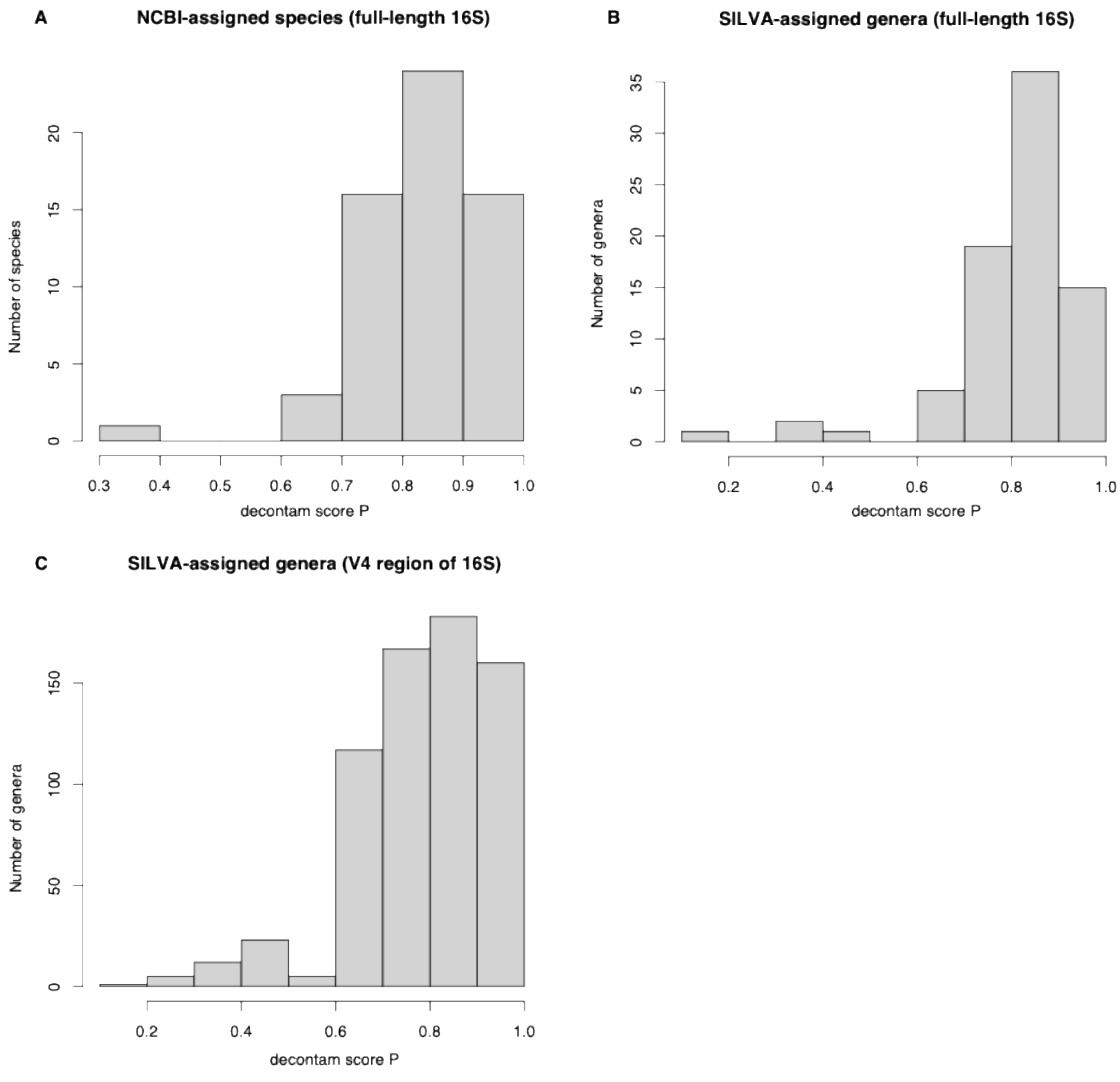

Supplementary Table 3. Summary of the number of reads produced by ONT (n=3) and Illumina (n=1) sequencing runs.

| Sequencing run ID | Total | Lowest per sample | Highest per sample | Average per sample | Median per sample | Number of genera detected |
|---|---|---|---|---|---|---|
| ONT Run 1 | 378742 | 9525 | 31514 | 21041.22 | 21938 | 75 |
| ONT Run 2 | 102580 | 2508 | 8789 | 5698.889 | 6099.5 | 75 |
| ONT Run 3 | 98616 | 2485 | 8427 | 5478.667 | 5826 | 75 |
| Illumina | 575425 | 3259 | 55136 | 26155.68 | 25901.5 | 20 |

Supplementary Figure 2. Comparison of actual and observed relative abundances of bacterial species found in the ZymoBIOMICS Microbial Community Standard, based on long-read full-length *16S* rRNA gene sequencing. Actual mock composition was provided by the manufacturer (Zymo Research, 2023). Composition of the Zymo Microbial Community Standard was consistent between sequencing runs and replicates, thus all replicates were pooled in the relative abundance plots.

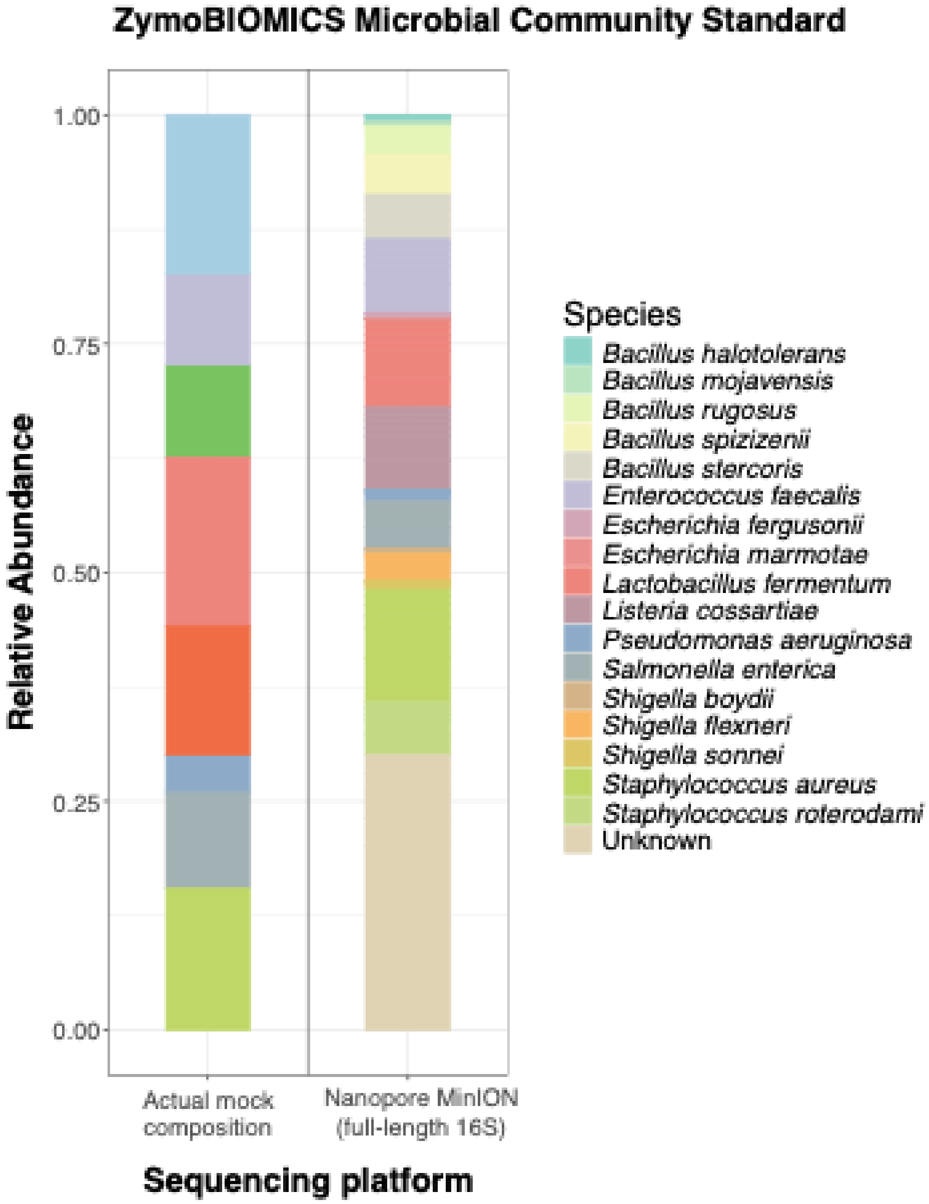

Supplementary Figure 3. Comparison of observed relative abundances of bacterial genera found within the ZymoBIOMICS Microbial Community Standard based on long-read full-length *16S* rRNA gene sequencing, assigned using the NCBI *16S/18S* and SILVA 138 *SSU* databases. Composition of the Zymo Microbial Community Standard was consistent between sequencing runs and replicates, thus all replicates were pooled in the relative abundance plots. SILVA curates the closely related genera *Escherichia* and *Shigella* together as the combined category *Escherichia-Shigella* while NCBI curates the two genera separately. These assignments were made consistent as *Escherichia-Shigella* and assigned the same green color.

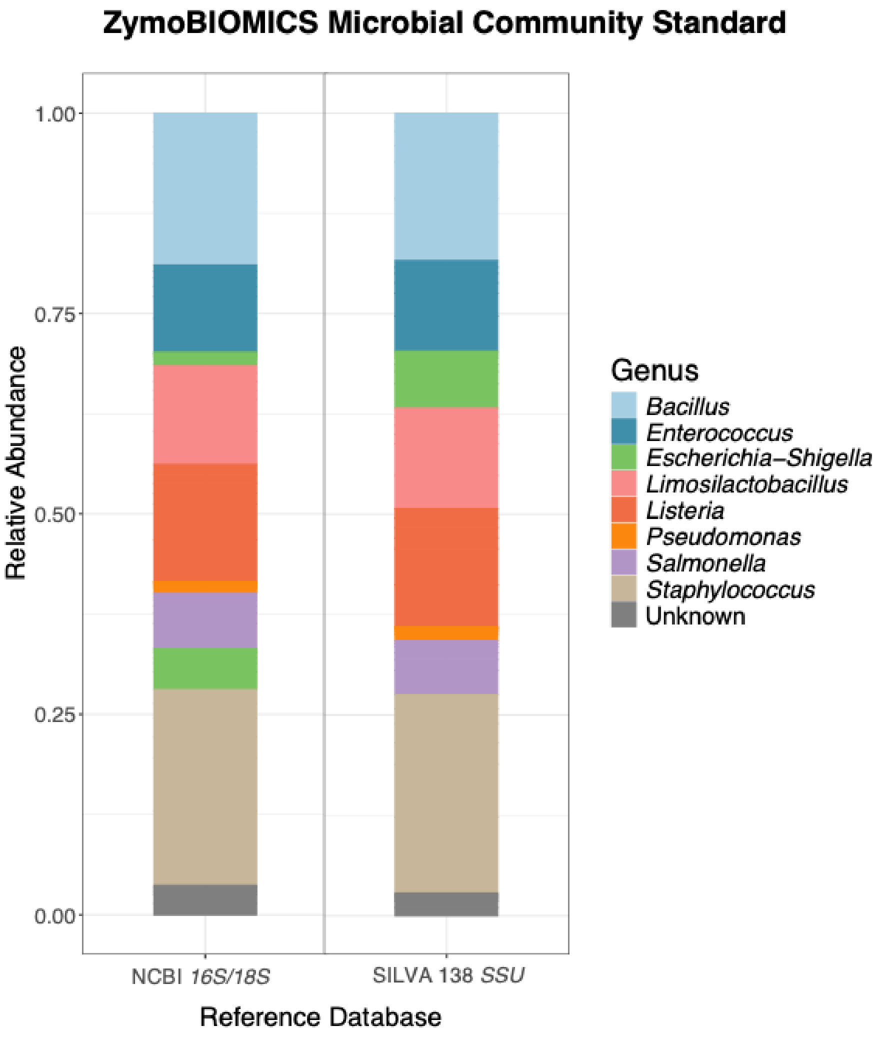
